# Multiple pathways of toxicity induced by *C9orf72* dipeptide repeat aggregates and G_4_C_2_ RNA in a cellular model

**DOI:** 10.1101/2020.09.14.297036

**Authors:** Frédéric Frottin, Manuela Pérez-Berlanga, F. Ulrich Hartl, Mark S. Hipp

**Affiliations:** Max Planck Institute of Biochemistry, Am Klopferspitz 18, 82152 Martinsried, Germany; Department of Quantitative Biomedicine, University of Zurich, Winterthurerstrasse 190, 8057 Zurich, Switzerland; Department of Biomedical Sciences of Cells and Systems, University Medical Center Groningen, University of Groningen, 9713 AV Groningen, The Netherlands; School of Medicine and Health Sciences, Carl von Ossietzky University Oldenburg, Oldenburg, Germany

## Abstract

The most frequent genetic cause of amyotrophic lateral sclerosis and frontotemporal dementia is a G_4_C_2_ repeat expansion in the *C9orf72* gene. This expansion gives rise to translation of aggregating dipeptide repeat (DPR) proteins, including poly-GA as the most abundant species. However, gain of toxic function effects have been attributed to either the DPRs or the pathological G_4_C_2_ RNA. Here we analyzed in a cellular model the relative toxicity of DPRs and RNA. Cytoplasmic poly-GA aggregates, generated in the absence of G_4_C_2_ RNA, interfered with nucleocytoplasmic protein transport, but had little effect on cell viability. In contrast, nuclear poly-GA was more toxic, impairing nucleolar protein quality control and protein biosynthesis. Production of the G_4_C_2_ RNA strongly reduced viability independent of DPR translation and caused pronounced inhibition of nuclear mRNA export and protein biogenesis. Thus, while the toxic effects of G_4_C_2_ RNA predominate, DPRs exert additive effects that may contribute to pathology.

## Introduction

Expansion of a GGGGCC hexanucleotide repeat (hereafter G_4_C_2_) within the first intron of the *C9orf72* gene is the most frequent genetic cause of amyotrophic lateral sclerosis (ALS) and frontotemporal dementia (FTD) (Renton et al., 2011). Mutant *C9orf72* in patients suffering from ALS/FTD can have more than a thousand G_4_C_2_ repeats, while healthy individuals possess usually less than twenty repeats (Gijselinck et al., 2016; Nordin et al., 2015). Transcripts with expanded G_4_C_2_ tracts are translated by repeat associated non-AUG (RAN) translation in all reading frames and in both strands, resulting in the synthesis of five different dipeptide repeat proteins (DPRs): poly-GA, poly-GR, poly-GP, poly-PR and poly-PA (Mori, Arzberger, et al., 2013; Mori, Weng, et al., 2013; Zu et al., 2013), all of which have been detected in patient brains (Mori, Arzberger, et al., 2013; Mori, Weng, et al., 2013; Zu et al., 2013). Poly-GA is the most abundant of the DPRs, followed by the other sense strand-encoded forms (poly-GP and poly-GR) (Mori, Weng, et al., 2013; Schludi et al., 2015). In patient brain and cellular models, DPRs accumulate in deposits that can be found in the nucleus and cytoplasm, including neurites (Ash et al., 2013; Gendron et al., 2013; Mori, Arzberger, et al., 2013; Mori, Weng, et al., 2013; Schludi et al., 2015; Zu et al., 2013). Poly-GA aggregates are localized mainly in the cytoplasm (Zhang et al., 2016), whereas arginine-containing DPRs (R-DPRs; polyGR and polyPR) accumulate in the nucleus, specifically the nucleoli (Kwon et al., 2014; May et al., 2014; Moens et al., 2019; Wen et al., 2014; White et al., 2019; Yamakawa et al., 2015; Zhang et al., 2014). However, in patients, poly-GR and poly-PR predominantly form cytoplasmic inclusions, with only a fraction of cells containing para-nucleolar inclusions that co-localize with silent DNA (Schludi et al., 2015). Interestingly, the less frequent intranuclear poly-GA inclusions in both cell models and patient brain are excluded from the nucleoli (Schludi et al., 2015).

Both loss- and gain-of-function mechanisms have been suggested to contribute to *C9orf72* pathology (reviewed in (Balendra & Isaacs, 2018)). Despite its location in a non-coding part of the gene, the G_4_C_2_ expansion can alter the expression level of the C9ORF72 protein (Rizzu et al., 2016; Waite et al., 2014). However, *C9orf72* knockout mouse models have failed to fully recapitulate ALS- or FTD-related neurodegenerative phenotypes, suggesting that loss of C9ORF72 protein is not the only contributor to pathology (Burberry et al., 2016; Burberry et al., 2020; Koppers et al., 2015; O’Rourke et al., 2016; Sudria-Lopez et al., 2016; Zhu et al., 2020). Toxic functions induced by this expansion have been studied in various cellular and animal models, and non-exclusive RNA- and protein-based mechanisms of toxicity have been proposed. However, the main contributor to gain of function toxicity in the disease remains to be defined.

mRNA containing the pathological G_4_C_2_ expansion has been shown to form stable G-quadruplexes in the nucleus that retain RNA binding proteins and induce splicing defects (Cooper-Knock et al., 2014; Donnelly et al., 2013; Gitler & Tsuiji, 2016; Haeusler et al., 2014; Sareen et al., 2013; Simon-Sanchez et al., 2012; Xu et al., 2013). R-DPRs have been additionally shown to interact with membrane-free phase separated compartments, such as the nucleolus, causing nucleolar stress and dysfunction of nucleolar quality control, as well as impairment of nucleocytoplasmic trafficking (Frottin et al., 2019; Hayes et al., 2020; Kwon et al., 2014; Lee et al., 2016; Mizielinska et al., 2017; Shi et al., 2017; Tao et al., 2015; Vanneste et al., 2019; White et al., 2019). Additionally, cytoplasmic poly-GA aggregates associate extensively with proteasomes, interfere with proteasome activity (Guo et al., 2018), and have been found to colocalize with p62, ubiquitin and several components of the proteasome machinery in patient brain (Al-Sarraj et al., 2011; Guo et al., 2018; May et al., 2014; Mori, Weng, et al., 2013). In mice expressing poly-GA, aggregates have been observed to sequester nuclear pore components (Zhang et al., 2016), thereby interfering with nucleocytoplasmic protein transport.

Differences between model systems as well as discrepancies in DNA constructs and repeat lengths utilized in various studies have precluded a clear mechanistic understanding of *C9orf72* toxicity pathways. Here we used G_4_C_2_ repeat-containing constructs as well as synthetic constructs encoding DPR proteins of comparable lengths without generating G_4_C_2_ RNA to analyze the relative contributions of RNA and DPR species to cellular pathology. Focusing on nuclear and cytoplasmic toxicity pathways, we found that cytoplasmic, but not nuclear poly-GA aggregates impaired nucleocytoplasmic transport. However, nuclear poly-GA was more toxic and interfered with nucleolar protein quality control and protein synthesis. The G_4_C_2_-containing RNA species induced a strong accumulation of poly-adenylated mRNA within the nucleus and dramatically inhibited protein biosynthesis. Thus, poly-GA protein and G_4_C_2_ RNA interfere with multiple cellular key pathways, with the RNA component exerting the major toxic effects limiting cell viability.

## Results

### Nuclear and cytoplasmic poly-GA aggregates differ in toxicity

Both G_4_C_2_ RNA and DPR proteins resulting from the pathological *C9orf72* expansion have been suggested to mediate gain of toxic function effects in various models (reviewed in (Balendra & Isaacs, 2018). To investigate the contribution of poly-GA proteins to cellular pathology, we engineered ATG-driven synthetic constructs expressing 65 GA repeats fused C-terminally to GFP (GA_65_-GFP). Notably, these constructs do not contain any G_4_C_2_ hexanucleotide motifs (Figure 1A and Figure 1-figure supplement 1A).

**Figure 1.**
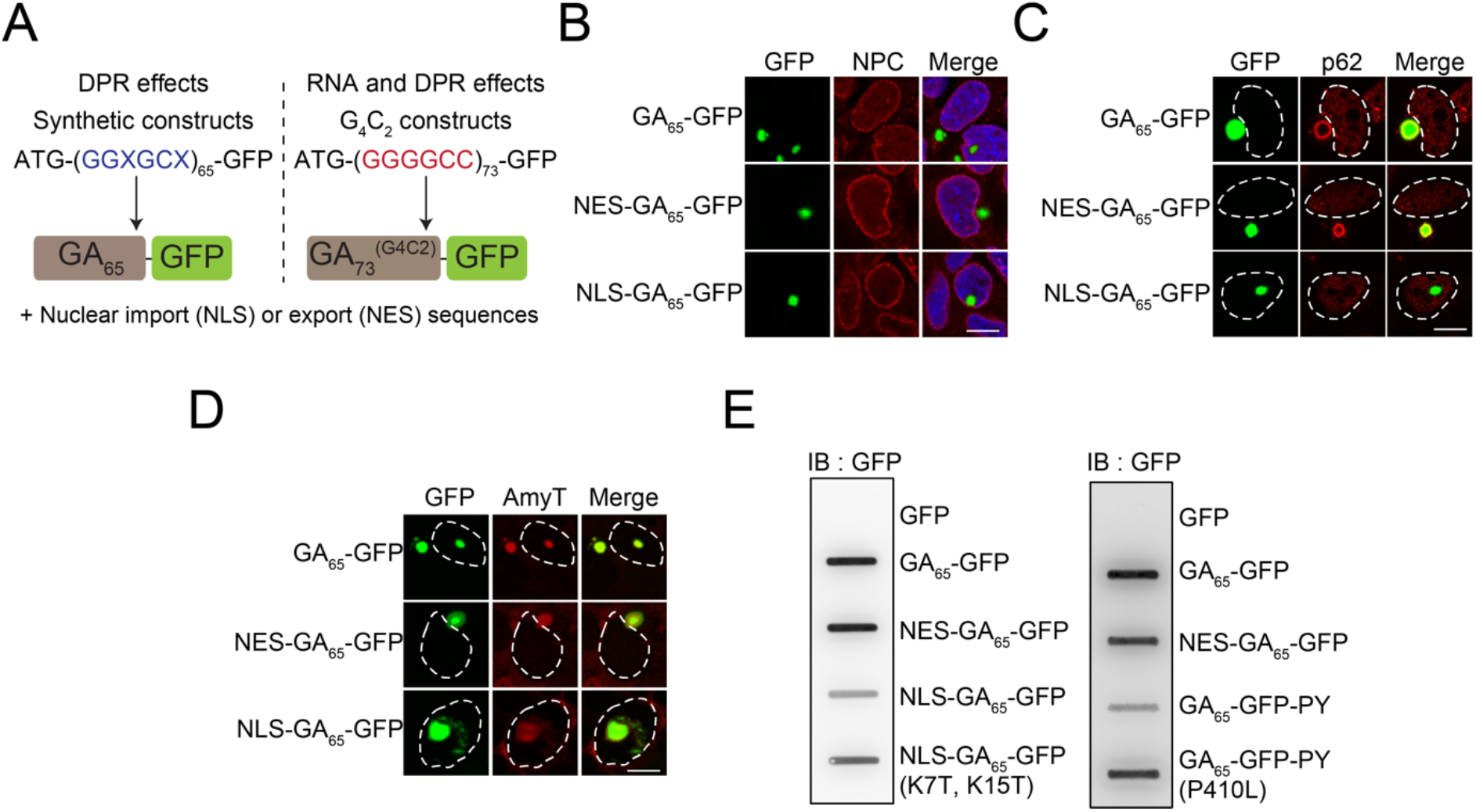
Cytoplasmic and nuclear poly-GA aggregates are amyloid-like but differ in solubility. (**A**) Schematic representation of the poly-GA encoding constructs used. Two different classes of poly-GA encoding constructs were used to study dipeptide repeat protein (DPR) mediated toxicity in presence or absence of RNA repeat regions: synthetic constructs that do not contain G_4_C_2_ motifs and are translated into GA_65_ (left), and constructs containing 73 G_4_C_2_ repeats that encode GA_73_ (right). Both classes contained ATG start codons and were fused in frame to GFP. N-terminal nuclear import (NLS) or export sequences (NES) were present when indicated. (**B**) The indicated constructs were transfected into HEK293T cells. Antibodies against nuclear pore complexes (NPC, red) were used to detect the nuclear membrane, poly-GA was visualized by GFP fluorescence (green), nuclei were counterstained with DAPI. (**C**) NES-GA_65_-GFP aggregates co-localize with p62. The indicated constructs were transfected into HEK293T cells. p62 (red) was detected by immunofluorescence and poly-GA was visualized by GFP fluorescence (green). White dashed lines delineate the nucleus based on DAPI staining. (**D**) Cytoplasmic and nuclear aggregates can be stained with an amyloid specific dye. The indicated constructs were transfected into HEK293T cells, and stained with AmyTracker (AmyT, red). Poly-GA DPRs were visualized by GFP fluorescence (green). White dashed lines delineate the nucleus based on DAPI staining. (**E**) Cytoplasmic GA aggregates are SDS insoluble. The indicated constructs were transfected into HEK293T cells. GFP was expressed as a soluble control protein. Cells were lysed and analyzed for SDS insoluble poly-GA aggregates by filter retardation assay. GFP antibody was used for detection. Scale bars represent 10 μm.

Expression of GA_65_-GFP in HEK293T cells resulted in the formation of bright fluorescent inclusions in most transfected cells. The inclusions were mostly cytoplasmic, except for a small fraction of cells with nuclear foci (Figure 1B). This phenotype is in agreement with reports on the localization of poly-GA aggregates in patient brain, where poly-GA aggregates are also observed in the neuronal cytoplasm and nucleus (Mori, Weng, et al., 2013; Schludi et al., 2015). Using engineered β-sheet proteins, we have previously reported that otherwise identical aggregation-prone proteins display distinct toxic properties when targeted to different cellular compartments (Frottin et al., 2019; Vincenz-Donnelly et al., 2018; Woerner et al., 2016). To test whether this is also the case for poly-GA, we generated compartment-specific variants of the poly-GA proteins. We restricted the expression of poly-GA to the cytoplasm by adding a nuclear export signal (NES-GA_65_-GFP) or targeted the protein to the nucleus using a double SV40 nuclear localization signal (NLS-GA_65_-GFP). NES-GA_65_-GFP accumulated in the cytoplasm and formed inclusions similar to those of GA_65_-GFP (Figure 1B). Directing the protein to the nucleus resulted in an increased number of cells with nuclear aggregates (Figure 1B). However, a number of NLS-GA_65_-GFP expressing cells also contained cytoplasmic inclusions, suggesting that aggregate formation in these cells occurred before the transport of the poly-GA proteins into the nucleus.

Poly-GA forms p62 positive inclusions in the cytoplasm of neurons (Guo et al., 2018). We were able to replicate this phenotype in our cellular system and observed p62-positive poly-GA inclusions of GA_65_-GFP and NES-GA_65_-GFP (Figure 1C) (May et al., 2014; Mori, Weng, et al., 2013). Both nuclear and cytoplasmic poly-GA inclusions were stained throughout with AmyT, a small amyloid-specific dye (Figure 1D). The aggregates were also recognized by the anti-amyloid antibody (OC) (Figure 1-figure supplement 1B), which recognizes generic epitopes common to amyloid fibrils and fibrillar oligomers (Kayed et al., 2007). However, while the nuclear aggregates stained homogenously with OC, cytoplasmic GA_65_-GFP inclusions showed only a peripheral reaction (Figure 1-figure supplement 1B). The differential antibody accessibility of the inclusion core suggests that the nuclear and cytoplasmic DPR aggregates, though both amyloidogenic, differ in structural properties such as packing density.

Analysis of the solubility of the nuclear and cytoplasmic GA_65_-GFP aggregates supported this interpretation. NES-GA_65_-GFP was retained in a filter retardation assay in the presence of SDS, while most NLS-GA_65_-GFP passed through the filter (Figure 1E). The difference in detergent solubility was not due to the presence of the NLS sequence, since similar results were obtained using the FUS-derived C-terminal nuclear localization signal (PY) (Gal et al., 2011) (Figure 1E). Moreover, disabling of the nuclear targeting sequences by point mutations resulted in the reappearance of SDS insoluble aggregates (Figure 1E). Thus, despite being amyloid-like, nuclear and cytoplasmic poly-GA aggregates have different physico-chemical properties.

The granular component (GC) of the nucleolus has recently been shown to function as a protein quality control compartment (Frottin et al., 2019). Interestingly, the nuclear aggregates of NLS-GA_65_-GFP altered the localization of the GC marker protein nucleophosmin (NPM1), while cytosolic NES-GA_65_-GFP had no such effect (Figure 2A). Nuclear GA_65_-GFP aggregates induced the dislocation of NPM1 from the GC phase of the nucleolus, with accumulation of NPM1 at the aggregate periphery (Figure 2A). However, the nuclear GA_65_-GFP deposits did not alter the distribution of the RNA polymerase I subunit RPA40, a marker of the fibrillar center of the nucleolus, the site of rRNA synthesis (Figure 2B). The NPM1-containing GC phase is responsible for pre-ribosome particle assembly and has been shown to accommodate proteins that have misfolded upon stress. These proteins enter the GC phase and are maintained in a state competent for refolding and repartitioning to the nucleoplasm upon recovery from stress (Frottin et al., 2019). In contrast to poly-GA, R-DPRs enter the GC phase of the nucleolus and convert it from liquid-like to a more hardened state, thereby impairing its quality control function (Frottin et al., 2019; Lee et al., 2016). To test whether nuclear poly-GA aggregates also affect nucleolar quality control, we expressed the synthetic poly-GA constructs in HEK293T cells together with nuclear firefly luciferase fused to the red fluorescent protein mScarlet (NLS-LS), a metastable protein that enters the GC phase upon stress-induced misfolding (Frottin et al., 2019). In control cells, NLS-LS accumulated in the nucleolus upon heat stress and largely repartitioned to the nucleoplasm within 2 h of recovery (Figure 2C, Figure 2-figure supplement 1A). A similar result was obtained upon expression of cytoplasmic NES-GA_65_-GFP. However, in the presence of aggregates of NLS-GA_65_-GFP, NLS-LS failed to efficiently repartition to the nucleoplasm (Figure 2C, Figure 2-figure supplement 1A), indicating that the nuclear poly-GA aggregates compromise nucleolar protein quality control.

**Figure 2.**
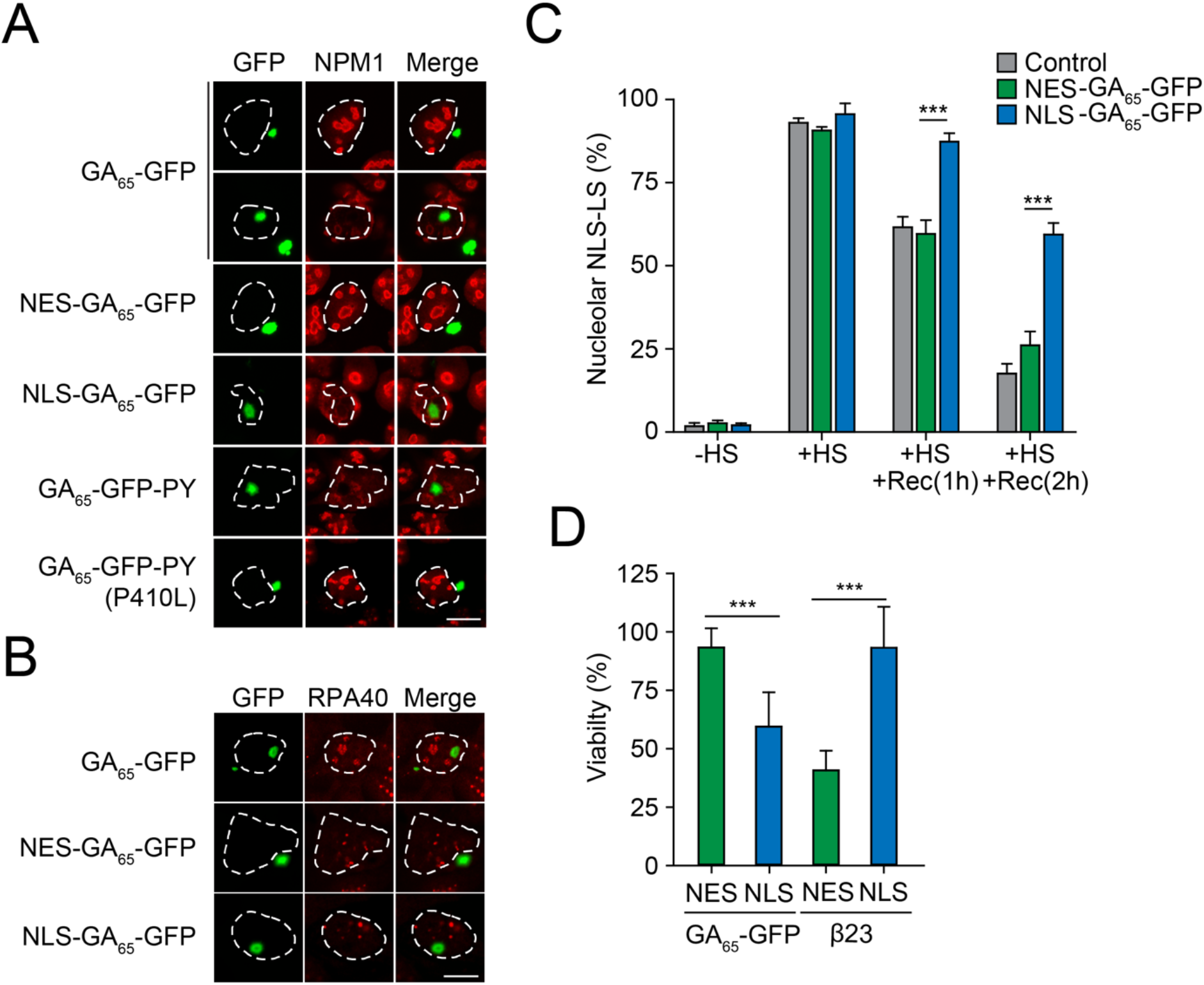
Nuclear poly-GA aggregates compromise nucleolar integrity and impair cell viability. (**A**) Nuclear poly-GA aggregates alter nucleophosmin (NPM1) localization. The indicated constructs were transfected into HEK293T cells. Cells were fixed and stained with anti-NPM1 antibodies (red). Poly-GA was visualized by GFP fluorescence (green). (**B**) Nuclear poly-GA aggregates are not nucleolar. The indicated constructs were transfected into HEK293T cells, followed by staining with antibodies against DNA-directed RNA polymerases I and III subunit RPAC1 (RPA40) (red). Poly-GA was visualized by GFP fluorescence (green). (**C**) Nuclear poly-GA aggregates disrupt nucleolar protein quality control. HEK293T cells were co-transfected with NLS-firefly luciferase fused to mScarlet (NLS-LS) and the indicated poly-GA constructs or GFP as a control. Cells were maintained at 37°C (−HS) or subjected to heat stress 43°C (+HS) for 2 h of heat stress and recovery (+HS +Rec) for 1 h and 2 h. Cells with nucleolar NLS-LS were counted and the results plotted as percentage of transfected cells. Data are shown as mean + SD (n = 3). *p*-Value of two-sided t-test is displayed (*** *p* ≤ 0.001). Representative immunofluorescence images are shown in Figure 2–figure supplement 1A. (**D**) Nuclear poly-GA is toxic. HEK293T cells were transfected with the indicated constructs and MTT cell viability assays were performed 4 days after transfection. Data were normalized to cells transfected with empty vector. Data are shown as means + SD (n ≥ 3);*p*-Values of two-sided t-test are shown (\*\*\**p* ≤ 0.001). White dashed lines delineate the nucleus based on DAPI staining. Scale bars represent 10 μm.

We next measured the viability of HEK293T cells expressing poly-GA in cytosol or nucleus using the MTT assay. While the expression of NES-GA_65_-GFP did not cause toxicity, cell viability was significantly reduced upon expression of NLS-GA_65_-GFP (Figure 2D). This effect was reproduced in cells transfected with a plasmid coding for GA_65_ from an alternative degenerated and G_4_C_2_-free DNA sequence (GA_65_(2)) (Figure 1-figure supplement 1A, Figure 2-figure supplement 1B), further excluding G_4_C_2_ mRNA as a cause of toxicity. Decreased viability was also observed when GA_65_-GFP was targeted to the nucleus via the alternative PY localization signal (Figure 2-figure supplement 1C). Importantly, both NES-GA_65_-GFP and NLS-GA_65_-GFP were expressed at levels comparable to GA_65_-GFP (Figure 2-figure supplement 1D). Point mutations in the NLS or PY targeting sequence prevented accumulation of GA_65_-GFP in the nucleus and restored viability (Figure 2-figure supplement 1C). Together these results indicate that the difference in toxicity between nuclear and cytoplasmic poly-GA is caused by compartment-specific properties of the aggregates independent of their targeting sequences and mRNA.

### Cytoplasmic GA_65_-GFP aggregates interfere with nuclear transport

We have previously shown that artificial β-sheet proteins, when aggregating in the cytoplasm, sequester nuclear transport factors and thereby interfere with transport of proteins and mRNA across the nuclear envelope (Woerner et al., 2016). Similar observations were made for the aggregates of various disease proteins including poly-GA and R-DPRs (Boeynaems et al., 2016; Chou et al., 2018; Eftekharzadeh et al., 2018; Freibaum et al., 2015; Gasset-Rosa et al., 2017; Grima et al., 2017; Jovicic et al., 2015; Khosravi et al., 2017; Kramer et al., 2018; Solomon et al., 2018; Zhang et al., 2015; Zhang et al., 2016). To test which role the localization of the poly-GA proteins plays in this process, we expressed NES-GA_65_-GFP or NLS-GA_65_-GFP together with the reporter protein shuttle-mApple (S-mApple). This reporter protein contains both nuclear import and export signals and consequently shuttles between nucleus and cytoplasm. At steady state, S-mApple localized mainly to the cytoplasm, but accumulated within minutes in the nucleus upon inhibition of nuclear export with leptomycin B (LMB) (Wolff et al., 1997) (Figure 3).

**Figure 3.**
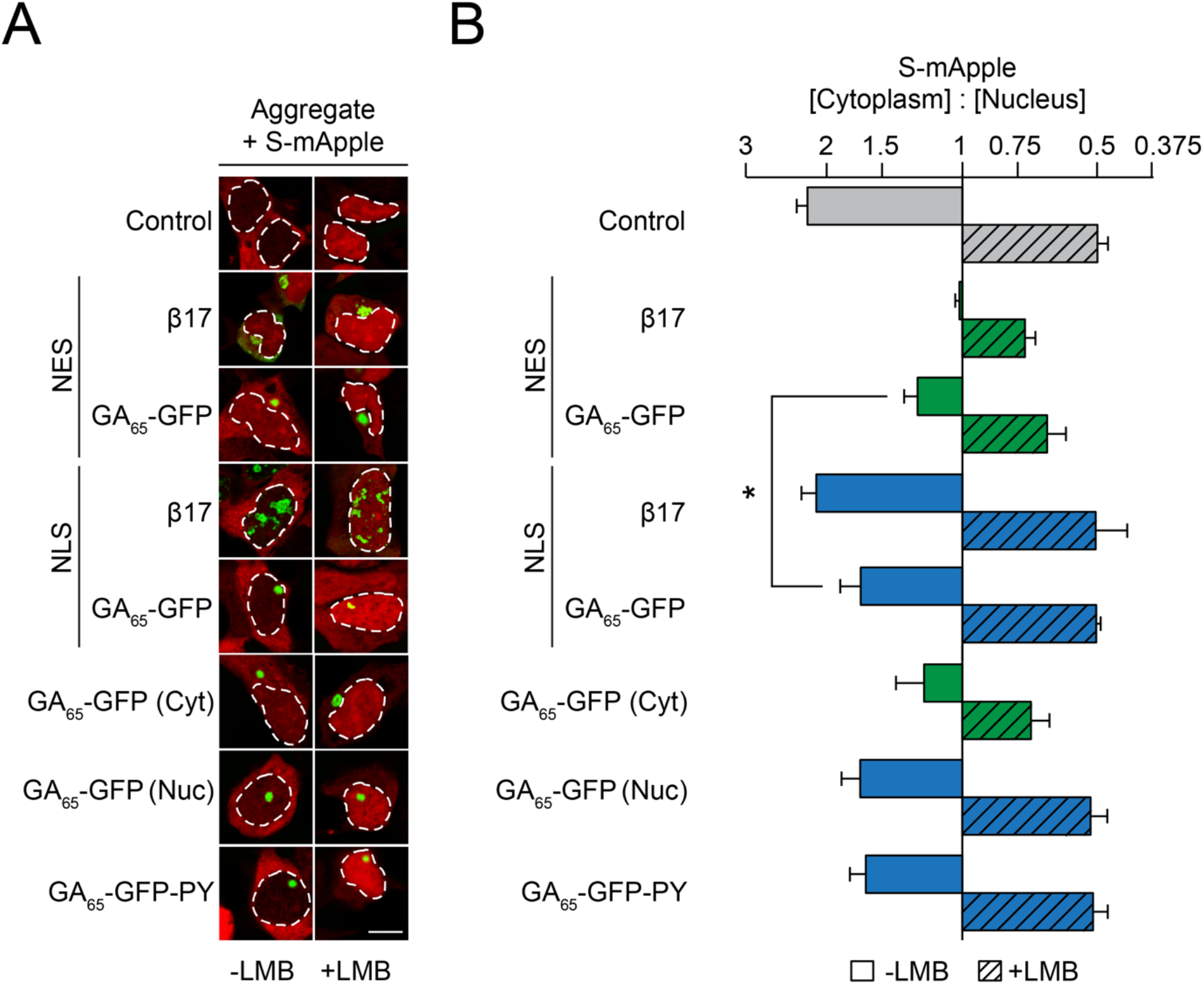
Cytoplasmic poly-GA aggregates impair nucleocytoplasmic protein transport. (**A**) Cytoplasmic poly-GA aggregates alter nuclear transport of a shuttling reporter protein. HEK293T cells were co-transfected with S-mApple (red) and either empty vector (Control), NES-β17, NES-GA_65_-GFP, NLS-β17, NLS-GA_65_-GFP, GA_65_-GFP or GA_65_-GFP-PY (green). Leptomycin B (LMB; 10 ng/ml) was added for 15 min when indicated. White dashed lines delineate nuclei based on DAPI staining. Scale bar represents 10 μm. (**B**) Quantification of S-mApple distribution from data in (A). The x-axis shows the enrichment of S-mApple concentration in the cytoplasm relative to the nucleus. Cells transfected with GA_65_-GFP were further analyzed and divided into cells with cytoplasmic (Cyt) or nuclear (Nuc) aggregates. Data are means + SD, n = 3 independent experiments, >70 cells were analzyed per condition. * *p* ≤ 0.05 from two-sided t-test.

As previously described, S-mApple was retained within the nucleus upon expression of the cytoplasmic β-sheet protein NES-β17 (Figure 3), indicative of inhibition of nuclear protein export (Woerner et al., 2016). Expression of NES-GA_65_-GFP also impaired S-mApple export from the nucleus (-LMB), but to a lesser extent than NES-β17 (Figure 3). A mild impairment of nuclear protein import by NES-GA_65_-GFP was also observed, as measured upon inhibition of export with LMB (Figure 3). As for NLS-β17, nuclear poly-GA aggregates had only a weak effect on protein export (Figure 3). Similarly, cells containing nuclear aggregates did not show a significant change in S-mApple import into the nucleus (Figure 3). These findings were replicated in cells displaying cytoplasmic or nuclear poly-GA aggregates of untargeted GA_65_-GFP (Figure 3).

We next monitored the nuclear translocation of p65, a subunit of the NF-κB complex, upon stimulation by the cytokine TNFα. In control cells, p65 is largely cytoplasmic and enters the nucleus upon treatment with TNFα (Figure 4A and B). Cells containing NES-GA_65_-GFP aggregates displayed a potent inhibition of p65 translocation (Figure 4A and B). A similar effect was seen with cytoplasmic aggregates of polyQ-expanded Huntingtin-exon 1 (Htt96Q) as a positive control (Figure 4A and B) (Woerner et al., 2016). Cells containing cytoplasmic aggregates of untargeted GA_65_-GFP also displayed reduced p65 translocation (Figure 4A and B). The observed translocation impairment was independent of an alteration of p65 phosphorylation and degradation of the inhibitor of nuclear factor κB (IκB) (Figure 4C). In contrast, nuclear poly-GA aggregates showed only a limited effect on p65 translocation (Figure 4A and B).

**Figure 4.**
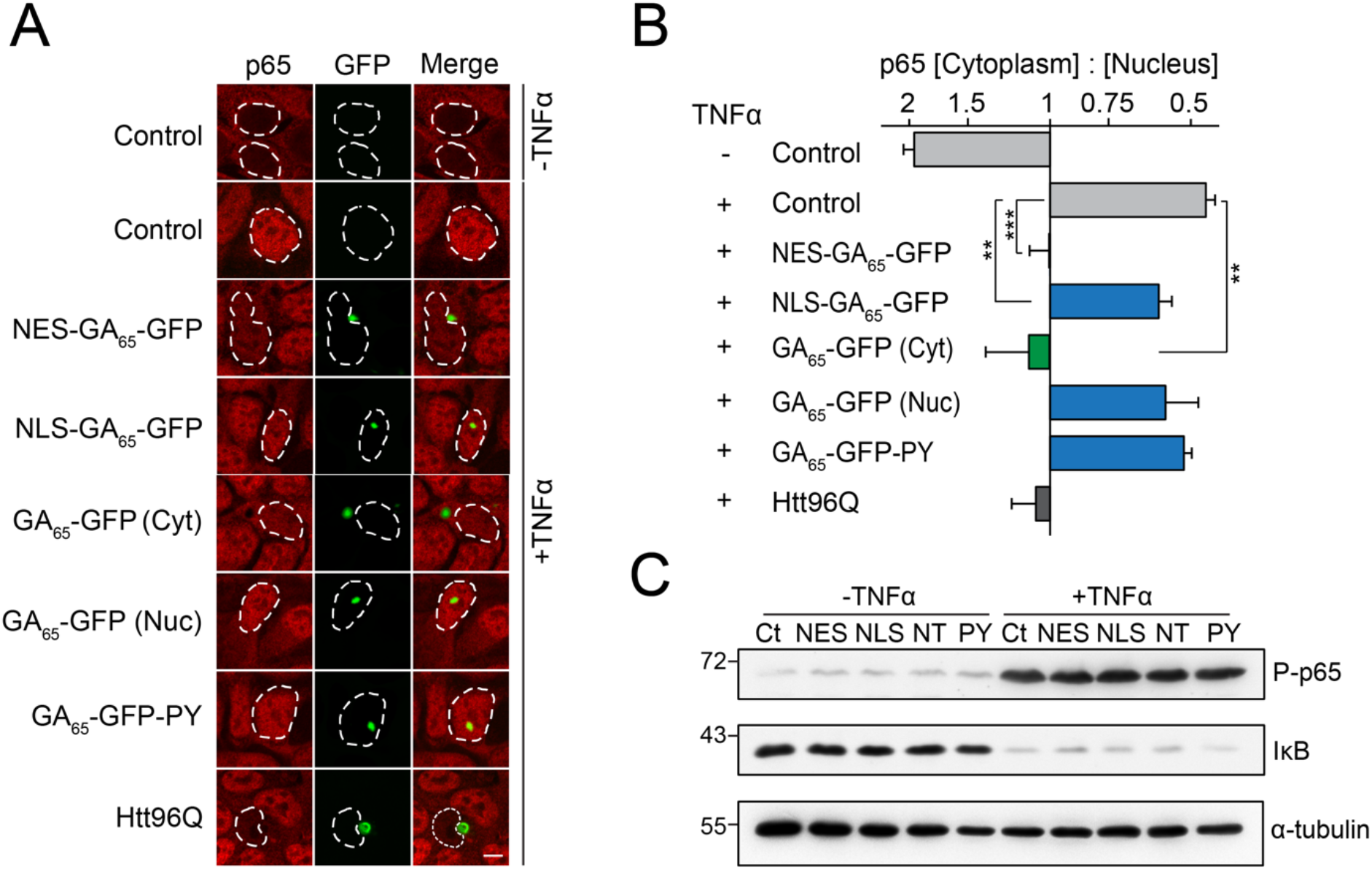
Cytoplasmic poly-GA aggregates inhibit nuclear import of p65. (**A**) Cytoplasmic poly-GA aggregates inhibit p65 nuclear translocation. HEK293T cells were transfected with empty vector (Control), NES-GA_65_-GFP, NLS-GA_65_-GFP, GA_65_-GFP, GA_65_-GFP-PY or Htt96Q-GFP (Htt96Q) (green) and analyzed for NF-κB p65 localization (red) with and without TNFα treatment (30 min). White dashed lines delineate nuclei based on DAPI staining. Scale bar represents 10 μm. (**B**) Quantification of NF-κB p65 distribution from data in (A). The x-axis shows the enrichment of p65 in the cytoplasm relative to the nucleus. Data are means + SD (n = 3), >100 cells were analyzed per condition. ** *p* ≤ 0.01, *** *p* ≤ 0.001 from two-sided t-test. (**C**) Expression of poly-GA does not alter the degradation of IκB and phosphorylation of p65. HEK293T cells were transfected with the indicated constructs (Ct, Control; NES, NES-GA_65_-GFP; NLS, NLS-GA_65_-GFP; NT, GA_65_-GFP; PY, GA_65_-GFP-PY) and treated as described in (A). Levels of IκB and phosphorylated NF-κB p65 (P-p65) were analyzed by immunoblotting. α-tubulin served as loading control.

Cytoplasmic aggregates of β-sheet protein and other disease-linked proteins cause mislocalization and sequester nuclear pore complexes and importins (Woerner et al., 2016). Particularly, R-DPRs have also been shown to directly bind and interfere with cargo loading onto importin at the nuclear pore (Hayes et al., 2020). While cytoplasmic poly-GA aggregates had no apparent effect on the localization of the nuclear pore complex (Figure 1B), we found that cells with cytoplasmic GA_65_-GFP aggregates also frequently contained aggregate foci of importins α1 (KPNA2), α3 (KPNA4) and β1 (KPNB1), not co-localizing with the poly-GA inclusions (Figure 4-figure supplement 1). Additionally, importins accumulated at the periphery of poly-GA inclusions (Figure 4-figure supplement 1). Nuclear poly-GA aggregates had no effect on the distribution of these importins (Figure 4-figure supplement 1). Thus, similar to the artificial β-sheet proteins, poly-GA aggregates induce compartment-specific cellular defects and impair nucleocytoplasmic protein transport.

### G_4_C_2_ repeat mRNA causes nuclear mRNA retention and pronounced toxicity

An aberrant distribution of mRNA has been observed in mouse motor neuron-like cells expressing expanded G_4_C_2_ repeats and in *C9orf72* patient cortical neurons (Freibaum et al., 2015; Rossi et al., 2015). We used an oligo-dT probe to test whether cytoplasmic poly-GA aggregates, generated from constructs lacking G_4_C_2_, affect the cellular distribution of total mRNA. In control cells, mRNA was present throughout the cytoplasm and in small nuclear ribonucleic particles (Figure 5A) (Carter et al., 1991). Expression of NES-GA_65_-GFP had only a minor effect on cellular mRNA distribution (Figure 5A, B), in contrast to the expression of cytoplasmic β-protein (NES-β17), which resulted in pronounced nuclear mRNA retention (Woerner et al., 2016). Likewise, NLS-GA_65_-GFP only caused limited mRNA retention in the nucleus (Figure 5A, B). Thus, poly-GA aggregates interfere with the function of only a subset of nuclear transport factors.

**Figure 5.**
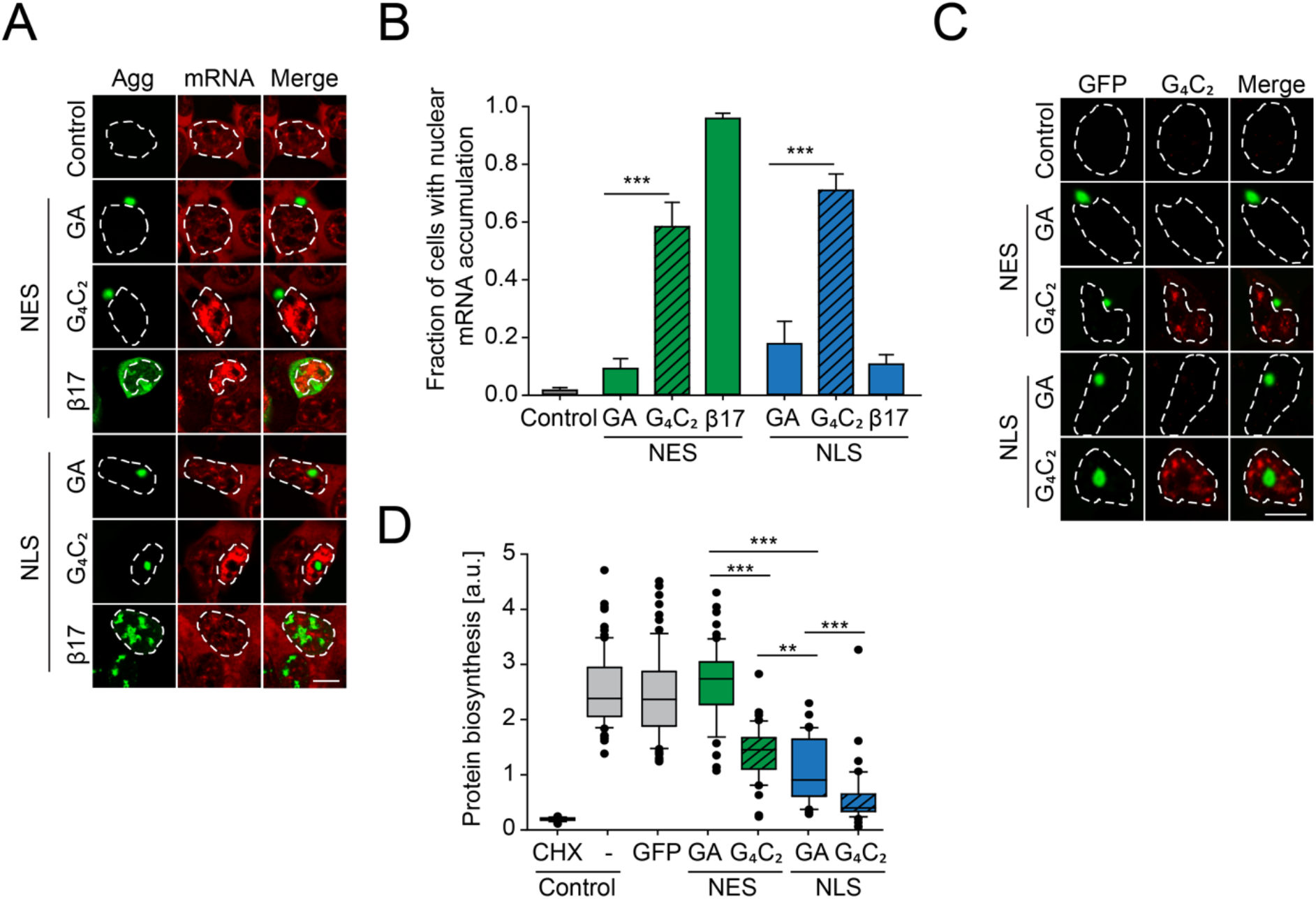
Protein biosynthesis defects correlate with retention of mRNA in the nucleus induced by G_4_C_2_ mRNA as well as presence of nuclear poly-GA aggregates. (**A**) G_4_C_2_ containing constructs induce strong nuclear mRNA accumulation. HEK293T cells were transfected with the indicated constructs: empty vector (Control); NES-GA_65_-GFP (NES-GA); NES-G_4_C_2_-GFP (NES-G_4_C_2_); NES-β17; NLS-GA_65_-GFP (NLS-GA); NLS-G_4_C_2_-GFP (NLS-G_4_C_2_); NLS-β17. PolyA-RNA was detected by fluorescence *in situ* hybridization using a poly-dT probe (red); protein aggregates (Agg) green. (**B**) Quantification of data in (A). The graph shows the fraction of cells with nuclear mRNA accumulation. Data are means + SD (n=3), *** *p* ≤ 0.001 from two-sided t-test. (**C**) G_4_C_2_ containing constructs induce the formation of G_4_C_2_ RNA foci. HEK293T cells were transfected with the indicated constructs:empty vector (Control); NES-GA_65_-GFP (NES-GA); NES-G_4_C_2_-GFP (NES-G_4_C_2_); NLS-GA_65_-GFP (NLS-GA); NLS-G_4_C_2_-GFP (NLS-G_4_C_2_). Cells were analyzed for GFP fluorescence (green) and C_4_G_2_ fluorescence by *in situ* hybridization (red). (**D**) Decreased protein biosynthesis caused by nuclear poly-GA and G_4_C_2_ mRNA. Protein biosynthesis was assayed in HEK293T cells transfected with the indicated constructs. Control cells were transfected with empty vector and treated with the translation inhibitor cycloheximide (CHX) when indicated. Boxplot of a representative experiment is shown. Center lines show the medians; box limits indicate the 25^th^ and 75^th^ percentiles; whiskers extend to the 10^th^ and 90^th^ percentiles, outliers are plotted as circles. Welch’s t-test was used to assess statistical significance (** *p* ≤ 0.01; *** *p* ≤ 0.001). The white dashed lines delineate the nucleus based on DAPI staining and the scale bars represent 10 μm.

Since the expression of poly-GA protein alone did not recapitulate the alterations of mRNA localization observed in *C9orf72* patient brain (Freibaum et al., 2015; Rossi et al., 2015), we next analyzed the effect of an ATG-driven poly-GA construct of 73 GA repeats encoded entirely by G_4_C_2_ motifs (G_4_C_2_)_73_ (Figure 1A and Figure 1-figure supplement 1A). Comparable to the synthetic GA_65_-GFP sequences, these constructs also generated poly-GA protein aggregates in the cytoplasm or nucleus, respectively (Figure 5A, C). However, both NES-(G_4_C_2_)_73_-GFP and NLS-(G_4_C_2_)_73_-GFP additionally generated G_4_C_2_ positive RNA foci in the nucleus, as observed by fluorescent in situ hybridization (FISH) using a C_4_G_2_ probe (Figure 5C). Cells expressing GA_65_-GFP or (G_4_C_2_)_73_-GFP without targeting sequence were analyzed as well, but only the G_4_C_2_ constructs showed mRNA accumulation in the nucleus in the majority of cells (Figure 5-figure supplement 1A). Furthermore, simultaneous visualization of G_4_C_2_ RNA puncta with the C_4_G_2_ probe, total mRNA using an oligo-dT probe and GA-GFP revealed that the presence of G_4_C_2_ RNA foci is associated with nuclear mRNA accumulation, independent of the presence of visible poly-GA protein aggregates (Figure 5-figure supplement 1B). Together these results indicate that G_4_C_2_ RNA, not poly-GA protein, mediates retention of mRNA in the nucleus.

The nuclear mRNA retention in cells expressing G_4_C_2_ constructs was accompanied by a marked reduction of protein synthesis as measured by the incorporation of a puromycin derivative into newly translated proteins (Slomnicki et al., 2016) (Figure 5D, Figure 5-figure supplement 1C). Interestingly, NLS-GA_65_-GFP expressing cells also displayed reduced protein synthesis, independently of G_4_C_2_ RNA. In contrast, NES-GA_65_-GFP had no inhibitory effect on protein synthesis (Figure 5D, Figure 5-figure supplement 1C). Notably, cells expressing NLS-(G_4_C_2_)_73_-GFP, accumulating both nuclear poly-GA protein and G_4_C_2_ RNA, were almost completely translation inactive, similar to control cells treated with the translation inhibitor cycloheximide (Figure 5D and Figure 5-figure supplement 1C). Thus, both nuclear poly-GA protein and G_4_C_2_ RNA appear to have additive inhibitory effects on protein biosynthesis. We next measured the viability of cells transiently transfected with either synthetic or G_4_C_2_-containing constructs. Independent of the presence of a NES or NLS targeting sequence, all G_4_C_2_ constructs markedly decreased cellular viability (Figure 6A), indicating that toxicity was mediated by the expanded G_4_C_2_ RNA. Moreover, expression of NLS-(G_4_C_2_)_73_-GFP was more toxic than NLS-GA_65_-GFP (Figure 6A), although the poly-GA levels of the constructs were comparable (Figure 6-figure supplement 1A).

**Figure 6.**
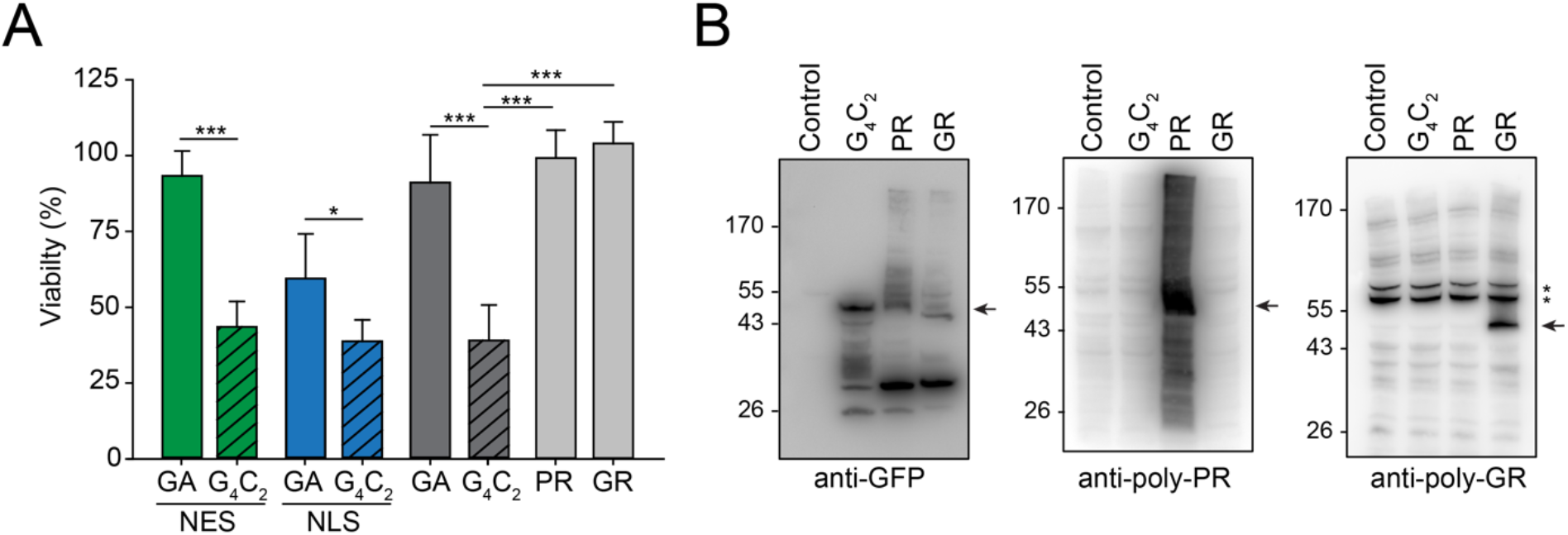
Production of G_4_C_2_ mRNA strongly decreases cellular viability. (**A**) G_4_C_2_ mRNA induces strong toxicity. HEK293T cells were transfected with the indicated constructs: G_4_C_2_-GFP (G_4_C_2_), NES-G_4_C_2_-GFP (NES-G_4_C_2_), NLS-G_4_C_2_-GFP (NLS-G_4_C_2_), GA_65_-GFP (GA), NES-GA_65_-GFP (NES-GA), NLS-GA_65_-GFP (NLS-GA), PR73-GFP (PR), or GR73-GFP (GR). MTT cell viability assays were performed 4 days after transfection. Data were normalized to cells transfected with empty vector. Data are means + SD (n ≥ 3). Part of this data is also shown in Figure 2D. \**p* ≤ 0.05; \*\*\**p* ≤ 0.001 from two-sided t-test. (**B**) (G_4_C_2_)_73_-GFP does not produce detectable amounts of arginine containing DPRs. HEK293T cells were transfected with the indicated constructs: empty vector (control), (G_4_C_2_)_73_-GFP (G_4_C_2_), PR73-GFP (PR), GR73-GFP (GR). Immunoblot analysis was then performed against GFP (left), poly-PR (center), and poly-GR (right). A representative result of three biological repeats is shown. The arrows indicate the main band of the respective DPRs and * indicate non-specific bands recognized by the anti-GR antibody.

Long sequences of repeated G_4_C_2_ motifs can produce a series of different DPRs by RAN translation (Ash et al., 2013; Gendron et al., 2013; Mori, Arzberger, et al., 2013; Mori, Weng, et al., 2013; Zu et al., 2013). Since the R-DPRs (poly-PR and poly-GR) have been reported to be toxic in cellular models (Boeynaems et al., 2016; Freibaum et al., 2015; Lee et al., 2016; Shi et al., 2017; Tao et al., 2015; Zu et al., 2013), we investigated whether the G_4_C_2_ constructs produced R-DPRs at levels sufficient to explain the observed toxicity independently of G_4_C_2_ RNA effects. To this end we engineered synthetic sequences resulting in the ATG-driven synthesis of 73 GR or 73 PR repeats (GR73-GFP and PR73-GFP, respectively) without G_4_C_2_ repeats. GR73-GFP was located predominantly within the cytoplasm and the nucleolus of HEK293T cells, while PR73-GFP accumulated within the nucleus and nucleolus (Figure 6-figure supplement 1B), as reported previously (Frottin et al., 2019; Lee et al., 2016; May et al., 2014; White et al., 2019). While the expression of (G_4_C_2_)_73_-GFP dramatically reduced cellular viability, the R-DPRs were not measurably cytotoxic (Figure 6A), consistent with an earlier report for constructs with longer repeat lengths (May et al., 2014). We used specific anti-GR and anti-PR antibodies to determine the relative accumulation of the R-DPRs in (G_4_C_2_)_73_-GFP expressing cells in comparison to cells expressing GR73-GFP and PR73-GFP. The antibodies recognized specific signals in cells transfected with the respective R-DPR constructs, but failed to detect R-DPR protein in cells expressing (G_4_C_2_)_73_-GFP (Figure 6B), indicating that production of R-DPR protein from the G_4_C_2_ constructs was very inefficient. Given that PR73-GFP and GR73-GFP were produced in detectable quantities from the synthetic constructs without inducing toxicity, we conclude that the pronounced toxicity observed upon expression of (G_4_C_2_)_73_-GFP cannot be explained by RAN translation of R-DPR but is due to nuclear mRNA retention mediated by the G_4_C_2_ repeat RNA.

## Discussion

We have employed a cellular model to differentiate possible mechanisms of toxicity exerted by expansion of the G_4_C_2_ hexanucleotide tract within the *C9orf72* locus, the most frequent genetic cause of ALS and FTD. Our results demonstrated that the G_4_C_2_ expansion causes cellular toxicity in a manner dependent on both aggregates of G_4_C_2_-encoded DPR proteins and the G_4_C_2_ repeat mRNA. These findings suggest a multiple hit model of additive, but mechanistically independent, effects of proteotoxicity and RNA-mediated toxicity, with the latter being the predominant toxic agent in the model system investigated.

Both DPRs and repeat mRNA interfered with different aspects of nucleocytoplasmic transport. The G_4_C_2_ RNA inhibited the export of mRNA from the nucleus, consistent with a dramatic impairment of protein synthesis that would be associated with strong neuronal toxicity. Notably, this effect was independent of the presence of DPR protein aggregates. Additionally, as has been reported for artificial β-sheet proteins and other disease-associated aggregation-prone proteins such as Tau, Huntingtin, FUS and TDP-43 (Chou et al., 2018; Dormann et al., 2010; Eftekharzadeh et al., 2018; Gasset-Rosa et al., 2017; Grima et al., 2017; Woerner et al., 2016), cytoplasmic poly-GA aggregates partially inhibited the transport of proteins across the nuclear pore. This observation is in agreement with the previously reported sequestration of transport factors by poly-GA (Khosravi et al., 2017; Zhang et al., 2016). Interestingly, other DPRs have also been reported to inhibit additional aspects of nucleocytoplasmic transport. Poly-PR can directly obstruct the central channel of the nuclear pore by binding to the FG domains of nuclear pore proteins (Shi et al., 2017), and R-DPRs can disrupt cargo loading onto karyopherins (Hayes et al., 2020).

While the effects of cytoplasmic poly-GA (NES-poly-GA) aggregates on nuclear transport were relatively well tolerated in our cell culture model, poly-GA aggregates within the nucleus (NLS-poly-GA) were associated with substantial proteotoxicity. Nuclear poly-GA formed aggregates at sites that were distinct from nucleoli, similar to the localization of poly-GA aggregates observed in patient brain (Schludi et al., 2015). However, formation of these inclusions altered the shape of nucleoli as a result of mislocalization and partial sequestration of the abundant GC protein NPM1. Nuclear poly-GA aggregates interfered with the recently described protein quality control function of the GC phase of the nucleolus (Frottin et al., 2019), as demonstrated using the metastable firefly luciferase as a model substrate. In cells containing nuclear poly-GA aggregates, misfolded luciferase failed to repartition from the nucleolus to the nucleoplasm during recovery from stress. This inhibitory effect was comparable to that previously observed for positively charged DPRs, such as poly-PR, which accumulate directly within the GC phase of the nucleolus (Frottin et al., 2019; Mizielinska et al., 2017; Tao et al., 2015). However, R-DPRs apparently do not accumulate in the nucleolus in patient brain (Schludi et al., 2015), and thus may rather interfere with nucleolar quality control by the mechanism described here for nuclear poly-GA (Frottin et al., 2019; Kwon et al., 2014; White et al., 2019).

Expression of poly-GA from G_4_C_2_ containing constructs resulted in substantially greater toxicity than expression from synthetic constructs, when similar poly-GA lengths and amounts were compared. The G_4_C_2_ hexanucleotide repeat is thought to exert toxic effects in part by forming higher order RNA assemblies in the nucleus which sequester RNA-binding proteins (Almeida et al., 2013; Cooper-Knock et al., 2015; Cooper-Knock et al., 2014; Donnelly et al., 2013; Haeusler et al., 2014; Mori, Lammich, et al., 2013; Reddy et al., 2013; Rossi et al., 2015; Sareen et al., 2013; Xu et al., 2013). We found that G_4_C_2_ constructs always resulted in strong inhibition of protein synthesis, even when coding for cytoplasmic poly-GA (NES-GA_65_), which itself alone did not impair protein biogenesis when produced from a synthetic construct. However, because expression constructs based on G_4_C_2_ repeats also generate the different RAN translation DPR products, a clear distinction between RNA and DPR toxicity in previous studies had been difficult. We therefore compared not only the relative toxicity of poly-GA and G_4_C_2_ constructs, but also measured the toxicity of poly-PR and poly-GR constructs in the absence of G_4_C_2_ repeat RNA. Notably, expression of these protein-only constructs did not induce overt toxicity, even when expressed at levels much higher than those generated by RAN translation of the G_4_C_2_ repeat. These findings allowed us to unequivocally attribute the major component of toxicity associated with G_4_C_2_ constructs to the production of the pathological G_4_C_2_ RNA sequence. Indeed, DPRs may be undetectable in the brain regions most affected by neurodegeneration in *C9orf72* patients (Schludi et al., 2015). We note however, that nuclear poly-GA also interfered with protein biosynthesis, presumably by impairing the nucleolar function in ribosome biogenesis, and thus could enhance the toxic effects of G_4_C_2_ repeat RNA.

In summary, our results indicate that the G_4_C_2_ expansion in *C9orf72* interferes with multiple nuclear functions, culminating in an inhibition of protein biogenesis, an outcome that would be especially harmful to neuronal cells. These dominant toxic effects might be further aggravated by a loss of function of the endogenous C9ORF72 protein, which is thought to play a role in cellular quality control (Boivin et al., 2020; Sellier et al., 2016; Sullivan et al., 2016; Yang et al., 2016; Zhu et al., 2020). Further research on the relative contribution of the different toxic mechanisms will be important in developing therapeutic strategies.

## Materials and Methods

### Cell culture, transfection, and cell treatments

Human embryonic kidney 293T (HEK293T) cells were obtained from ATCC and maintained in Dulbecco’s modified Eagle’s medium (DMEM) (Biochrom KG) supplemented with 10% fetal bovine serum (Gibco), 100 U/ml penicillin and 100 μg/ml streptomycin sulfate (Gibco), 2 mM L-glutamine (Gibco) and 5 μg/ml Plasmocin (InvivoGen). For heat stress (HS) and recovery experiments, cells were either maintained at 37 °C (-HS), or placed in a 43 °C (+HS) incubator for the indicated duration, or subjected to heat stress and then transferred back to 37 °C for recovery (+HS +Rec). Transient transfections were performed by electroporation with the GenePulser XCell System (Bio-Rad) or with Lipofectamine 2000 (Invitrogen) according to the manufacturer’s instructions. For assessing nuclear import of p65, transfected cells were treated with 20 ng/ml recombinant human TNFα (Jena Biosciences) for 30 min. For translation inhibition, cycloheximide (CHX, Sigma-Aldrich) was dissolved in phosphate buffered saline (PBS) and applied at a final concentration of 1 mM.

### Plasmids

Degenerated sequences encoding 65 GA^ggxgcx^ repeats preceded by a start codon (ATG) and flanked by NheI and BamHI restriction sites were chemically synthesized by GeneArt Gene Synthesis (Invitrogen). The GC content of sequence 1 is 77.2%, while sequence 2 contains 81.5% GC (Figure 1-figure supplement 1A). Both GA_65_ sequences were cloned into pcDNA3.1-myc/His A plasmids in frame with GFP at the C-terminus. An N-terminal nuclear export signal (NES) or an N-terminal double SV40 nuclear localization signal (NLS) were inserted by site-directed mutagenesis. The alternative FUS-derived nuclear localization signal (PY) was inserted C-terminally. Point mutations were introduced by site-directed mutagenesis. Similarly, degenerated sequences encoding PR73 and GR73 were generated by GeneArt Gene Synthesis (Invitrogen) and fused to the N-terminus of GFP. All sequences contained an ATG start codon. Sequences encoding (G_4_C_2_)_73_ repeats preceded by a start codon (ATG) were generated as previously described by primer hybridization (Guo et al., 2018). Similarly, the (G_4_C_2_)_73_ construct was subcloned in place of the degenerated GA_65_ sequence in the same NES/NLS-tagged GFP-containing vector. A GFP-only construct was generated by deletion of the GA from the same vector. NLS-LS has been previously described (Frottin et al., 2019). The plasmid encoding for S-mApple was generated from mApple-N1 (Addgene plasmid # 54567) a kind gift from Michael Davidson (Shaner et al., 2008). mApple was then cloned between BamHI and XbaI to replace GFP in a previously described plasmid encoding shuttle GFP (Woerner et al., 2016). c-myc-NES-β17, c-myc-NLS-β17 and Htt96Q plasmids have been previously described (Woerner et al., 2016). All relevant plasmid regions were verified by sequencing.

### Antibodies and dyes

The following primary antibodies were used in this study: nuclear pore complex proteins (Mab414), Abcam (24609); NPM1, Invitrogen (32-5200); NF-κB p65 (D14E12), Cell Signaling Technology (#8242); c-Myc-Cy3 (9E10), Sigma (C6594); amyloid fibrils, OC, Millipore (AB2286); GFP, Roche (11814460001); α-tubulin, Sigma Aldrich (T6199); RPA40, Santa Cruz (sc-374443); IκBα (L35A5), Cell Signaling Technology (#4814); phospho-NF-κB p65 (Ser536) (93H1), Cell Signaling Technology (#3033); KPNA2, Abcam (ab70160); KPNA4, Abcam (ab84735); KPNB1, Abcam (ab2811); GAPDH, Millipore (MAB374); poly-PR, Proteintech (23979-1-AP); poly-GR (5A2), Millipore (MABN778). The following secondary antibodies were used: Mouse IgG-Alexa488, Cell signaling Technology (#4408); Mouse IgG-Alexa555, Cell signaling Technology (#4408); Rabbit IgG-Alexa555, Cell signaling Technology (#4408); Mouse IgG-Alexa488, Invitrogen (21200); Mouse IgG-Alexa 633, Invitrogen (A21053); anti-mouse IgG-Peroxidase, Sigma Aldrich (A4416); anti-rabbit IgG-Peroxidase, Sigma Aldrich (A9169). The amyloid dye AmyTracker 680 (AmyT; Ebba Biotech AB) was used as previously described (Frottin et al., 2019). Briefly, cells were fixed in 4% paraformaldehyde in PBS (GIBCO) for 20 min, washed with PBS, permeabilized with Triton X-100 0.1 % for 5 min. AmyT was used at 1:500 dilution and incubated with the samples for 1 h at room temperature. Nuclei were counterstained with 4’, 6-Diamidine-2’-phenylindole dihydrochloride (DAPI, Molecular Probes).

### Immunofluorescence and image acquisition

Cells were grown on poly-L-lysine coated coverslips (Neuvitro). Cells were fixed with 4% paraformaldehyde, permeabilized with 0.1% Triton X-100, and blocked with 1% bovine serum albumin in PBS. Primary antibodies were applied in blocking buffer supplemented with 0.1% Triton X-100 and incubated overnight at 4°C. Appropriate fluorescent secondary antibodies at a dilution of 1:500 were applied for 60 min at room temperature. Nuclei were counterstained with DAPI before mounting samples with fluorescence-compatible mounting medium (DAKO).

Confocal microscopy was performed at MPIB Imaging Facility (Martinsried, Germany) on a ZEISS (Jena, Germany) LSM780 confocal laser scanning microscope equipped with a ZEISS Plan-APO 63x/NA1.46 oil immersion objective. In case of multi-fluorescence samples, a single-stained control sample was used to adjust emission and detection configuration to minimize spectral bleed-through. Images of cells with inclusions for co-localization studies were subjected to linear unmixing with spectra obtained from the single-stained samples using ZEN software. When fluorescence intensities were directly compared, acquisition settings and processing were kept identical. Images were analyzed with ImageJ (Rasband, W.S., National Institutes of Health, USA) and assembled in Adobe Photoshop CC (Adobe Systems Incorporated, Release 19.1.5).

### Cell viability assay

HEK293T cells were transfected by electroporation as previously described (Woerner et al., 2016). In brief, cells were electroporated with 20 μg of plasmid in 0.4 cm-gap electroporation cuvettes (Bio-Rad). Cells were electroporated at 225 V, ∞ Ω, 950 μF exponential wave in a GenePulser XCell System (Bio-Rad). After electroporation, cells were plated in a 24-well plate in triplicates. MTT assays were performed 3 days after transfectrion. The growth medium was replaced with fresh medium containing 5 μg/ml thiazolyl blue tetrazolium bromide (Sigma) for 1 h. Formazan crystals were solubilized by addition of stop solution, containing 40% N,N-dimethylformamide (Sigma-Aldrich), 16% SDS (Sigma-Aldrich) and 2% (v/v) acetic acid (Sigma-Aldrich). Absorbance at 570 nm and 630 nm was then recorded. Alternatively, viability was measured using the CellTiter-Glo 2.0 Cell Viability Assay kit (Promega) in the same conditions.

### Determination of nuclear import/export with S-mApple

Cells were co-transfected with the indicated constructs and the reporter S-mApple. After 48 h, cells were treated with 10 ng/ml of the CRM1 inhibitor Leptomycin B (LMB, Sigma Aldrich) in DMSO for 15 min before cell fixation. Control cells received DMSO. The relative concentration of S-mApple in the cytoplasm and nucleus was quantified by measuring the fluorescence intensity ratio in cells from 3 independent experiments. Fluorescence intensities were determined using ImageJ.

### Protein biosynthesis assay by click-chemistry

Protein biosynthesis assays were carried out using the click-it plus O-propargyl-puromycin (OPP) protein synthesis assay (ThermoFisher Scientific) according to the manufacturer’s instructions. Cells were transfected for 24 h with the indicated construct before metabolic labelling and incubated with the OPP reagent for 30 min in normal growth conditions. As control, protein translation was inhibited with cycloheximide (CHX, Sigma-Aldrich) dissolved in phosphate buffer saline (PBS) and applied at a final concentration of 1 mM. Samples were then fixed, permeabilized and the click reaction performed as recommended by the provider. Samples were subsequently analyzed by confocal microscopy. The concentration of labelled proteins was quantified by measuring the mean fluorescence intensity in 100-250 cells using ImageJ. A representative experiment of three independent experiments is shown.

### Fluorescence in situ hybridization (FISH)

Visualization of mRNA and G_4_C_2_ RNA by FISH was carried out as previously described (Woerner et al., 2016). HEK293T cells were fixed in 4% formaldehyde for 10 min at room temperature and permeabilized with 0.1% Triton X-100, both in UltraPure SCC buffer (Thermo Fisher Scientific). After washing with SSC buffer and with FISH buffer (10% formamide in SCC buffer), the probes (either T30 or (C_4_G_2_)5) were hybridized in FISH buffer with 10% dextran sulphate (Sigma Aldrich) for 3 h at 42°C, followed by additional washes in FISH buffer. When required, immunostaining was performed in PBS-based buffers afterwards. The fraction of transfected cells displaying abnormal accumulation of mRNA (using poly-dT probe) within the nucleus was determined by confocal microscopy.

### Filter retardation assay

For the filter retardation assay (Scherzinger et al., 1997; Wanker et al., 1999) cells were harvested 24 h after transfection with the indicated plasmids, lysed in radioimmunoprecipitation assay (RIPA) buffer (Thermo Scientific) and sonicated for 10 s. After incubation for 30 min, protein concentration was measured by Bradford assay (Bio-Rad), and equal amounts of lysates were filtered through a 0.2 μm pore size cellulose acetate membrane (GE Healthcare) and washed with lysis buffer. The membrane was subsequently immunoassayed with anti-GFP antibody. Antibody binding was detected using Luminata Forte Western HRP substrate (Millipore) and pictures were acquired with a LAS-3000 camera system (Fujifilm). AIDA (Raytest) software was used for analysis and quantitation.

### Statistics

Significance of differences between samples was determined using unpaired Student’s t test, ubless stated otherwise. Significance levels: (*) for *p* < 0.05, (**) for *p* < 0.01, (***) for p < 0.001.

## Author contributions

F.F. and M.P.B. designed and performed the experiments. F.F. supervised M.P.B..

F.F., M.S.H. and F.U.H. initiated and supervised the project, and wrote the paper.

## Funding

The research leading to these results has received funding from the European Commission under grant FP7 GA ERC-2012-SyG_318987–ToPAG. F.F. was supported by an EMBO Long Term Fellowship, M.P.B was supported by the Lehre@LMU Student Research Award Program of the Faculty of Biology at LMU Munich (sponsored by the Federal Ministry of Education and Research, funding nr. 01PL17016).

## Acknowledgements

We thank R. Klein, D. Edbauer, R. Körner and R. Sawarkar for helpful discussions. We acknowledge technical support by the MPIB Imaging facility.

## Competing Interests

No competing interests declared.

## Supplementary Information

**Figure 1–figure supplement 1.**
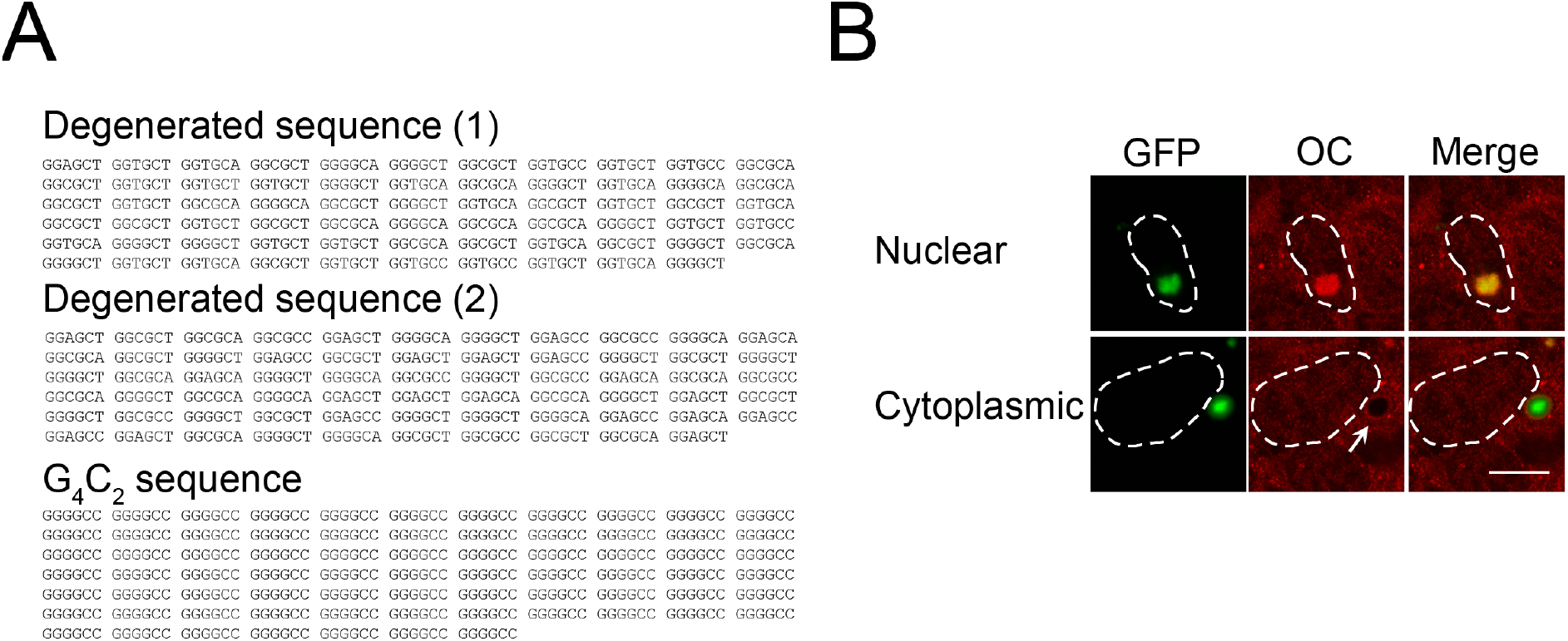
Cytoplasmic and nuclear poly-GA aggregates differ in their staining properties. (**A**) Sequences of the GA expressing constructs used in this study. Sequences (1) and (2) are synthetic degenerated sequences that do not contain G_4_C_2_. A third construct only contains G_4_C_2_ repeats and is translated into GA_73_. All three constructs contain an ATG start codon in frame with the GA coding sequence. (**B**) Cytoplasmic and nuclear aggregates can be stained with the OC antibody against amyloid fibrils. GA_65_-GFP was expressed in HEK293T cells, and immunofluorescence analysis was performed against amyloid fibrils (OC, red). Poly-GA was visualized by GFP fluorescence (green). White dashed lines delineate the nucleus based on DAPI staining. Scale bar represents 10 μm.

**Figure 2–figure supplement 1.**
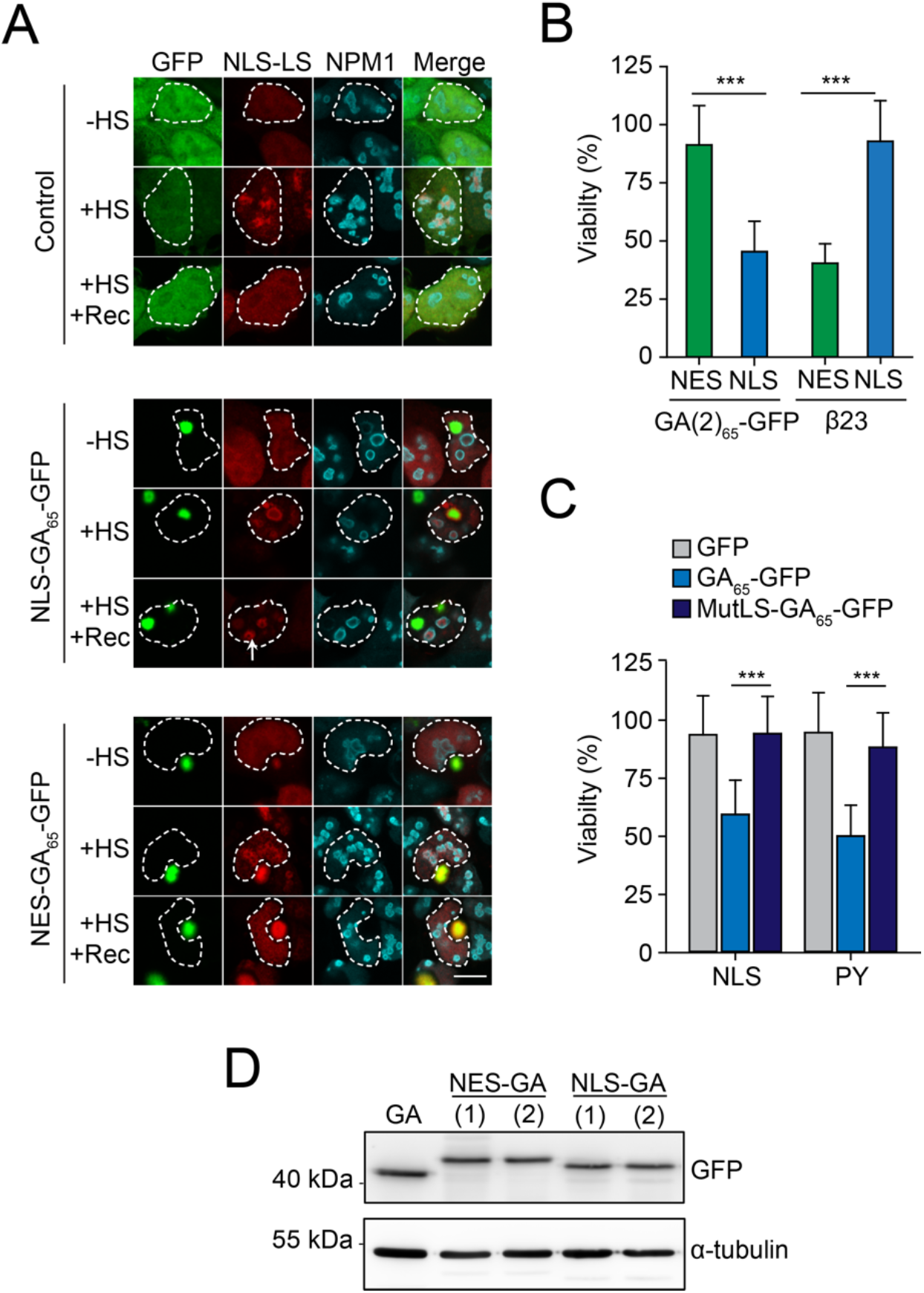
Nuclear poly-GA aggregates impair nucleolar protein quality control and cell viability. (**A**) Nuclear poly-GA aggregates disrupt nucleolar protein quality control. HEK293T cells were co-transfected with NLS-LS and the indicated poly-GA (NLS-GA_65_-GFP or NES-GA_65_-GFP) constructs or GFP as control. Cells were maintained at 37°C (−HS) or subjected to heat stress at 43°C for 2 h (+HS) and recovery (+HS +Rec) for 2 h. Samples were immunostained for NPM1 (cyan), NLS-LS and poly-GA proteins were visualized by mScarlet and GFP fluorescence, respectively. Merged panels are shown and a white dashed line delineates the nucleus based on DAPI staining. Scale bars represent 10 μm. Representative images of 3 independent experiments are shown. See Figure 2C for quantification. (**B**) NLS-GA_65_-GFP produced from an alternative degenerated and G_4_C_2_-free DNA sequence also decreases cellular viability. HEK293T cells were transfected with the indicated constructs and MTT cell viability assays performed 4 days after transfection. Data were normalized to cells transfected with empty vector. Data are means + SD (n ≥ 3) and two-sided t-test is shown (*** *p* ≤ 0.001). β23 data are from Figure 2D and are shown as control. (**C**) Disabling nuclear targeting sequence functionality of GA_65_-GFP rescues cellular viability. HEK293T cells were transfected with constructs encoding two different nuclear localization sequences (NLS or PY) fused to GFP, GA_65_-GFP or GA_65_-GFP with an inactivating mutation in the nuclear localization sequence (MutLS-GA_65_-GFP), followed by MTT cell viability assay 4 days after transfection. Data were normalized to cells transfected with empty vector. Data are shown as means + SD (n ≥ 3). *** *p* ≤ 0.001 by two-sided t-test. Part of this data is also shown in Figure 2D. (**D**) HEK293T cells were transfected with the indicated constructs (GA_65_-GFP; NES-GA_65_-GFP; NES-GA_65_(2)-GFP; NLS-GA_65_-GFP; and NLS-GA_65_(2)-GFP). Expression levels were determined by immunobloting with anti-GFP antibodies, α-tubulin served as loading control.

**Figure 4–figure supplement 1.**
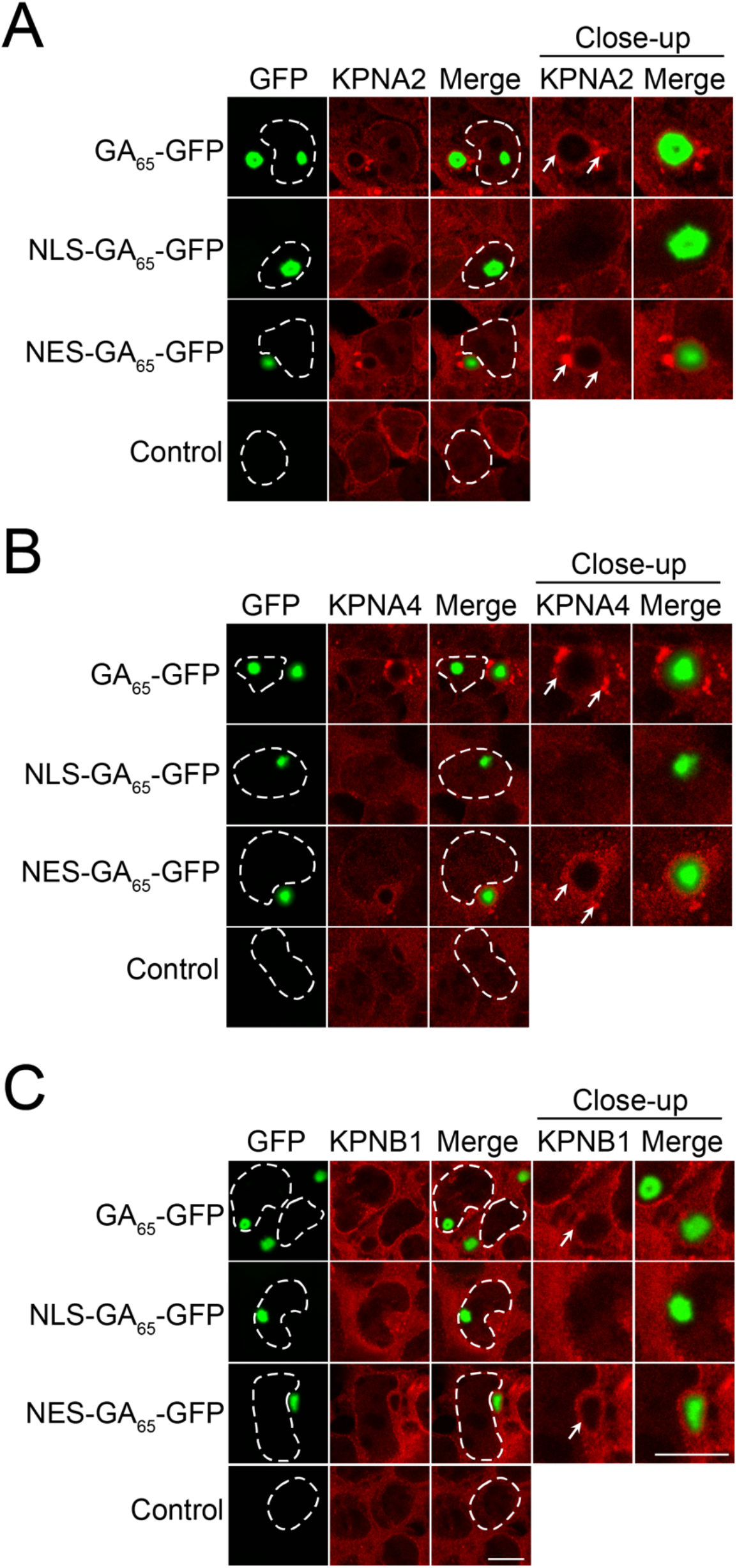
Importins form aggregates in cells containing cytoplasmic poly-GA aggregates and are partially sequestered. HEK293T cells were left untreated (Control) or were transfected with GA_65_-GFP, NES-GA_65_-GFP or NLS-GA_65_-GFP (green). Cells were then analyzed by immunofluorescence with antibodies against importin α-1 (KPNA2; (**A**)), importin α-3 (KPNA4; (**B**)) and importin β-1 (KPNB1; (**C**)) (red). Close-up views of inclusions are shown and arrows indicate importin aggregates and/or their sequestration at the periphery of the inclusion. The white dashed lines delineate nuclei based on DAPI staining. Scale bars are 10 μm.

**Figure 5–figure supplement 1.**
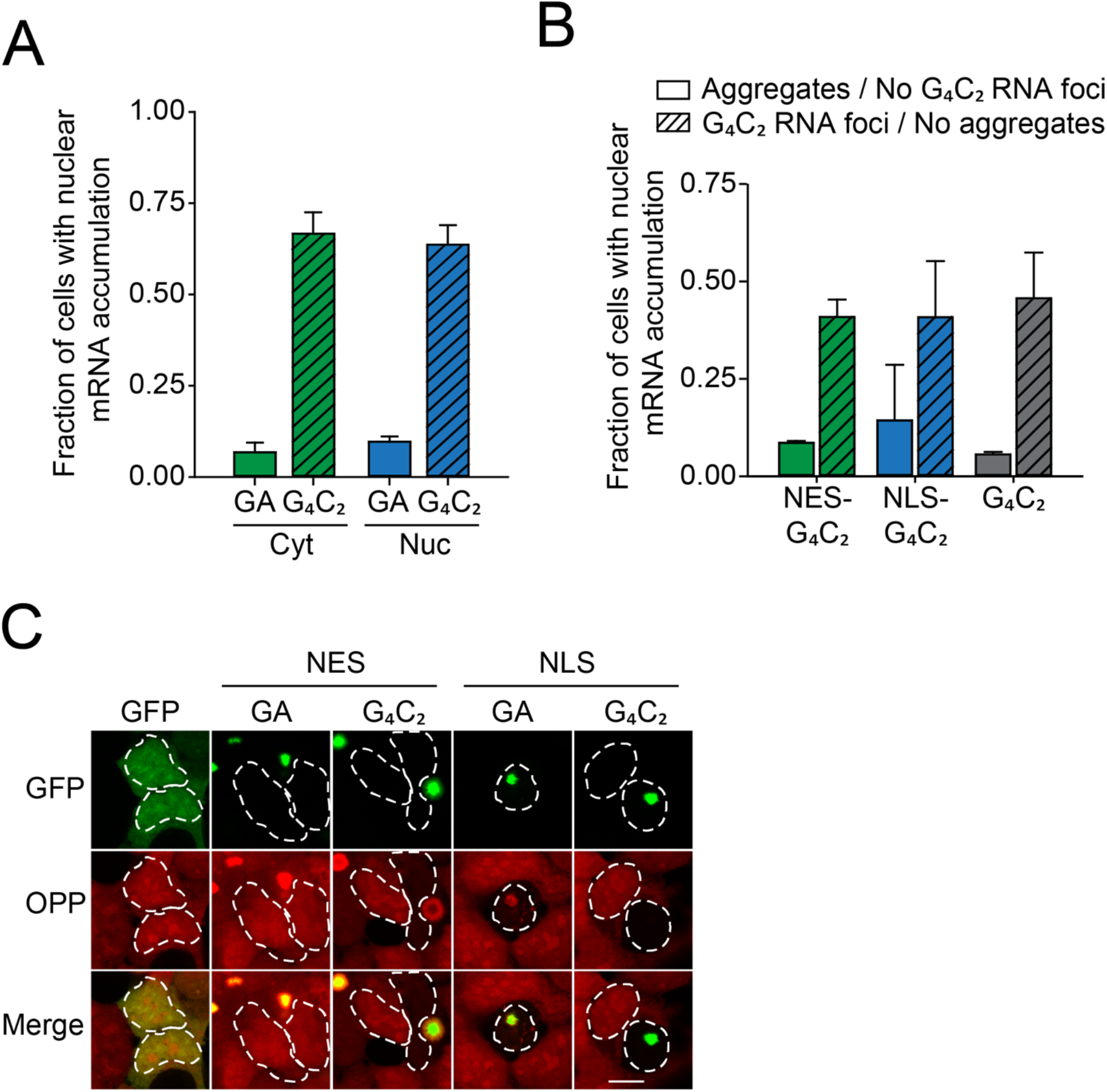
Expression of G_4_C_2_ containing constructs induces mRNA retention in the nucleus. (**A**) Presence of G_4_C_2_ rather than aggregate localization causes nuclear mRNA retention. HEK293T cells were transfected with either GA_65_-GFP (GA) or G_4_C_2_-GFP (G_4_C_2_) and analyzed for nuclear mRNA accumulation as in Figures 5A and B. Cells containing GA-GFP aggregates in the cytoplasm (Cyt) or nucleus (Nuc) were analyzed separately. Data are shown as means + SD (n = 3). (**B**) mRNA retention correlates with the presence of G_4_C_2_ positive nuclear foci. HEK293T cells were transfected with G_4_C_2_-GFP (G_4_C_2_), NES-G_4_C_2_-GFP (NES-G_4_C_2_) or NLS-G_4_C_2_-GFP (NLS-G_4_C_2_) and analyzed for mRNA nuclear accumulation as in Figure 5C. FISH was performed as in Figure 5C to detect G_4_C_2_ foci. Cells were categorized according to the presence of G_4_C_2_ positive foci as well as aggregates. Data are shown as means + SD (n = 3). (**C**) Decreased protein biosynthesis in presence of nuclear poly-GA and G_4_C_2_ mRNA. Newly synthesized proteins were labelled with O-propargyl-puromycin (OPP; red) in HEK293T cells transfected with the indicated constructs (green). Quantification of this experiment is shown in Figure 5D. The white dashed lines delineate the nucleus based on DAPI staining and the scale bar represents 10 μm.

**Figure 6–figure supplement 1.**
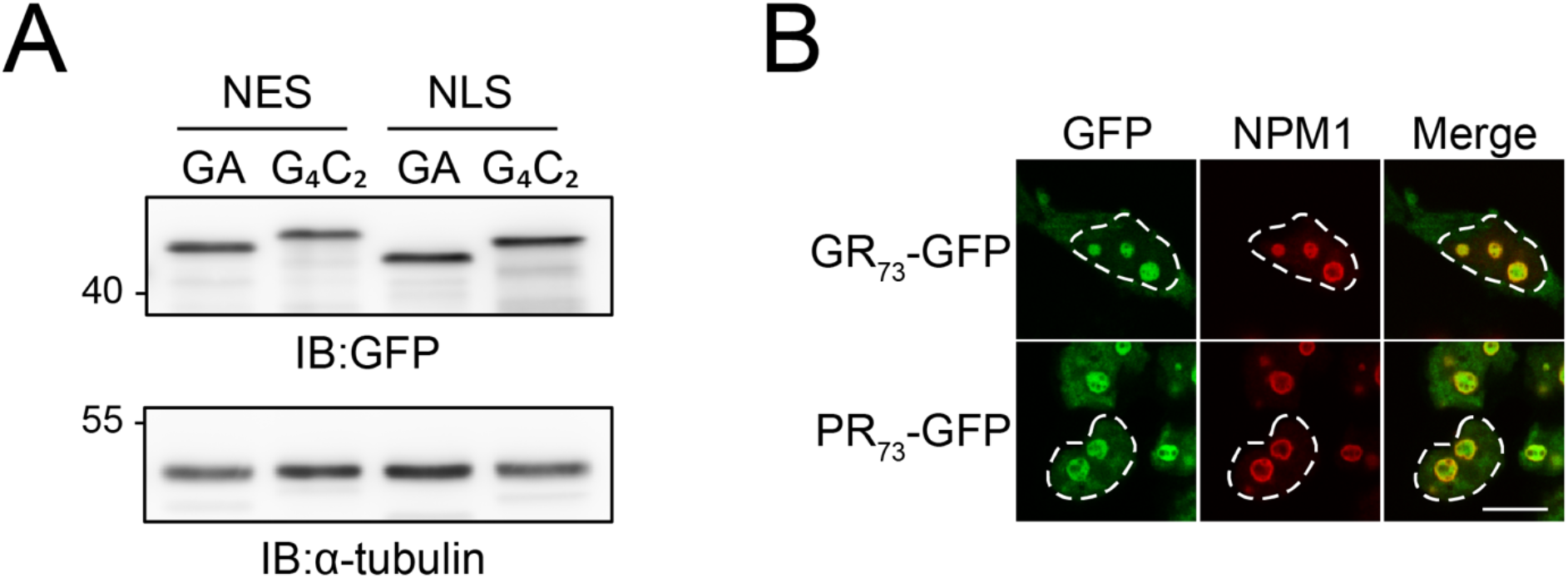
(**A**) Toxicity induced by G_4_C_2_ mRNA is not due to a higher poly-GA level. HEK293T cells were transfected with the indicated constructs: NES-GA_65_-GFP (NES-GA); NES-G_4_C_2_-GFP (NES-G_4_C_2_); NLS-GA_65_-GFP (NLS-GA); NLS-G_4_C_2_-GFP (NLS-G_4_C_2_). Poly-GA-GFP protein levels were analyzed by immunoblotting against GFP. α-tubulin served as loading control. A representative result of 3 independent experiments is shown. (**B**) Arginine containing DPRs accumulate in the nucleolus. HEK293T cells were transfected with the indicated constructs (GR73-GFP (GR); PR73-GFP (PR)). Cells were then stained with anti-nucleophosmin (NPM1, red). Expressed constructs are shown in green. The white dashed lines delineate the nucleus based on DAPI staining and the scale bar represents 10 μm.

